# Beyond traditional visual sleep scoring: massive feature extraction and unsupervised clustering of sleep time series

**DOI:** 10.1101/2021.09.08.458981

**Authors:** Nicolas Decat, Jasmine Walter, Zhao H. Koh, Piengkwan Sribanditmongkol, Ben D. Fulcher, Jennifer M. Windt, Thomas Andrillon, Naotsugu Tsuchiya

**Author notes:** These authors contributed equally to this work (Nicolas Decat, Jasmine Walter, Zhao H. Koh).

## Abstract

Sleep is classically measured with electrophysiological recordings, which are then scored based on guidelines tailored for the visual inspection of these recordings. As such, these rules reflect a limited range of features easily captured by the human eye and do not always reflect the physiological changes associated with sleep. Here we present a novel analysis framework that characterizes sleep using over 7700 time-series features from the *hctsa* software. We used clustering to categorize sleep epochs based on the similarity of their features, without relying on established scoring conventions. The resulting structure overlapped substantially with that defined by visual scoring and we report novel features that are highly discriminative of sleep stages. However, we also observed discrepancies as *hctsa* features unraveled distinctive properties within traditional sleep stages. Our framework lays the groundwork for a data-driven exploration of sleep and the identification of new signatures of sleep disorders and conscious sleep states.

## Introduction

### Background

Sleep is underpinned by a rich repertoire of biological processes. Overnight polysomnography (PSG) combining electroencephalography (EEG), electromyography (EMG) and electrooculography (EOG) shows that concerted neurophysiological events occur repeatedly across the night. The observation of cyclic patterns suggests that the brain goes through several states during sleep, each one being characterized by distinct patterns of underlying neural activity. A set of rules for classifying these states was first developed by Rechtschaffen and Kales (1). The authors designed these guidelines with awareness of their limitations. This scoring system was originally proposed as a reference point to facilitate comparisons of research findings among researchers and clinicians, expecting that the guidelines would be updated as research and technology evolved (2). Despite their original intentions, these guidelines are used to this day by the American Academy of Sleep Medicine (AASM) and have undergone only a few updates. The AASM visual scoring rules currently serve as the gold standard for the assessment of sleep physiology and the diagnosis of sleep disorders.

### Visual inspection of PSG data

The AASM sleep scoring manual provides guidelines for categorizing sleep and wake states in five discrete stages (Wake, N1, N2, N3, REM). These rules are based on the visual inspection of neurophysiological signals. Sleep technicians typically examine patterns of characteristic rhythms in the neural signals as well as specific physiological events to classify 30-second segments of sleep. For example, REM sleep is defined by the AASM as a stage of low amplitude, mixed frequency activity accompanied by low chin muscle tone and rapid eye movements. These rules, based on visually recognizable markers, are easy to describe and to standardize across laboratories, playing a significant role in the development of sleep research over the decades.

### Arbitrariness of scoring guidelines

Because the scoring guidelines have undergone only minimal revision since they were established, they mostly reflect the state of knowledge and practice from past decades (2). A typical example is the 30-second window rule, which was introduced at the time brain signals were recorded on 30 cm wide polygraph pages moving at 1 cm/s (3). Clearly, our current technology makes this obsolete. Regarding the PSG features that were selected for sleep scoring, some of them were chosen based on their visual clarity in preprocessed time series, with minimal contribution from modern data analysis methods. In fact, at the time the AASM sleep scoring guidelines were published, 26 of the 29 visual scoring rules were determined by consensus among human scorers and were proposed without a complete validation of their biological relevance to sleep (4, 5). For instance, the 75 μV amplitude criterion for slow waves is arbitrary (4) and is not suited to the assessment of sleep quality in older people, who often do not have delta oscillations this large (6, 7). Furthermore, some of the most distinctive features specified by the AASM, such as sleep spindles in N2 or rapid eye movements in REM, are neither completely specific nor completely necessary to the scoring of a stage, which hinders a reliable and accurate description of sleep data. The arbitrariness and incompleteness of current scoring rules and analytic approaches is a known but unresolved limitation in the field of sleep research and medicine.

### Inadequacy in characterizing sleep-stage transitions

One of the prominent problems that scorers often encounter regards epochs that are ambiguous and cannot be dealt with using AASM criteria. Such ambiguities arise around sleep-stage transitions, where the underlying biological states presumably change gradually. Hence, interscorer disagreement mainly occurs between adjacent stages (8). To cope with ambiguous epochs, the AASM recommends taking into account information from the neighboring epochs (9, 10). For example, a 30-second epoch with no characteristic EEG rhythms or transient events is scored as N2 if the epoch prior to or posterior to it shows sleep spindles. While this practice assumes continuity in the underlying sleep physiology and its dynamics, it is subject to highly variable criteria across scorers. Crucially, the analysis of sleep-stage transitions has revealed crucial insights in the diagnosis of several sleep disorders including narcolepsy (11-13), chronic fatigue syndrome (14), and insomnia (15). Furthermore, this ‘temporal smoothing’ process tends to underestimate transitional states, such as N1, which is not clearly identifiable visually but is associated with various processes like mental imagery (16) and memory processing (17). The contextual information rules, which are used to cope with the inability to visually identify and verbalize distinctive features, are not suited to the definition of sleep-stage transitions.

### Limited characterization of sleep time series

Despite all these known issues, traditional sleep scoring continues to rely on visually identifiable features in PSG signals. While raw EEG signals are noisy, non-stationary, and high dimensional, the traditional guidelines supplement visually identifiable features with only a few signal attributes, such as the dominant power in certain frequency bands. Thus, current guidelines do not fully capture the range of dynamical patterns that could underlie relevant changes in sleep physiology, and we might gain more understanding by characterizing sleep dynamics using the sophisticated range of modern time-series analysis algorithms.

### Purpose of this study

The goal of the present study is to organize sleep in a data-driven way, based on a wide range of their time-series properties. To achieve this goal, we first apply a massive feature extraction tool to sleep time series, using the highly comparative time-series analysis approach implemented in the *hctsa* software package which contains a diverse set of over 7700 time-series features (18, 19). *hctsa* encodes scientific algorithms for time-series analysis in the form of *features* that each summarize a structural property of a time series in the form of a real number. It includes methods that were developed in and applied to a wide range of research areas, including Fourier and wavelet transforms, entropy, self-correlation and predictability, nonlinear time-series analysis, and fractal scaling. This large set of analyses has never before been applied systematically to PSG data.

Based on the results of these *hctsa* time-series analyses, we sought to organize sleep (i.e., intra-sleep wakefulness, N1, N2, N3 and REM) independently of pre-defined scoring rules. To this end, we approached sleep scoring in an unsupervised way by clustering sleep time series based on the similarity of the dynamical properties captured by *hctsa*. Furthermore, we analyzed the data on an epoch-by-epoch basis to minimize subjective criteria related to the incorporation of contextual information. By taking this data-driven approach, which we refer to as feature-based clustering, our aim was not to replicate or improve AASM visual scoring, but to go beyond it. We examined this data-driven organization by comparing it to the traditional structure defined by AASM rules.

We present the analyses in three parts. First, we assess the extent to which our feature-based clusters correspond with visual (AASM) sleep scoring of the same epochs. Second, we examine cases in which feature-based clusters disagree with AASM labels to better understand what novel properties of sleep our approach can reveal. Third, we assess the ability of individual *hctsa* time-series features to distinguish AASM stages, identifying both traditional and novel features that promise new discoveries of the structure of sleep.

## Results

### Part 1. Correspondence between feature-based clustering and traditional sleep stage architecture

To objectively assess the correspondence between how sleep epochs are organized using (i) time-series features and (ii) AASM labels, we developed an automated procedure that combines unsupervised clustering, cluster matching, and cross validation (Figure 1; see Methods; *Feature-based clustering*). We applied *hctsa* to 12 full-night PSG datasets, each containing a preprocessed EEG, EOG, and EMG derivation. Through this process, the EEG, EOG and EMG time series of each 30-second epoch is converted into a single vector containing the output values produced by all *hctsa* features for this epoch (Figure 1a). The feature values which generated non-real numbers or errors were filtered out and the remaining valid values were normalized. After forming a balanced dataset (Figure 1b & c), we grouped 70% of the data (training set) into 5 clusters based on their *hctsa* feature values (Figure 1d). Using our sequential maximum matching algorithm, we assigned each unlabeled training cluster to one of the five AASM sleep stages (Figure 1e). We named the five training clusters C1, C2, C3, CR and CW, as those that most closely mapped onto N1, N2, N3, REM and wake, respectively. Then, we assigned the remaining 30% data (testing set) based on their distance to the training clusters (Figure 1f). We used the proportion of match between the AASM labels and the cluster labels of the test data (cross-validation) through 100 iterations to assess the correspondence between the two methods.

**Figure 1.**
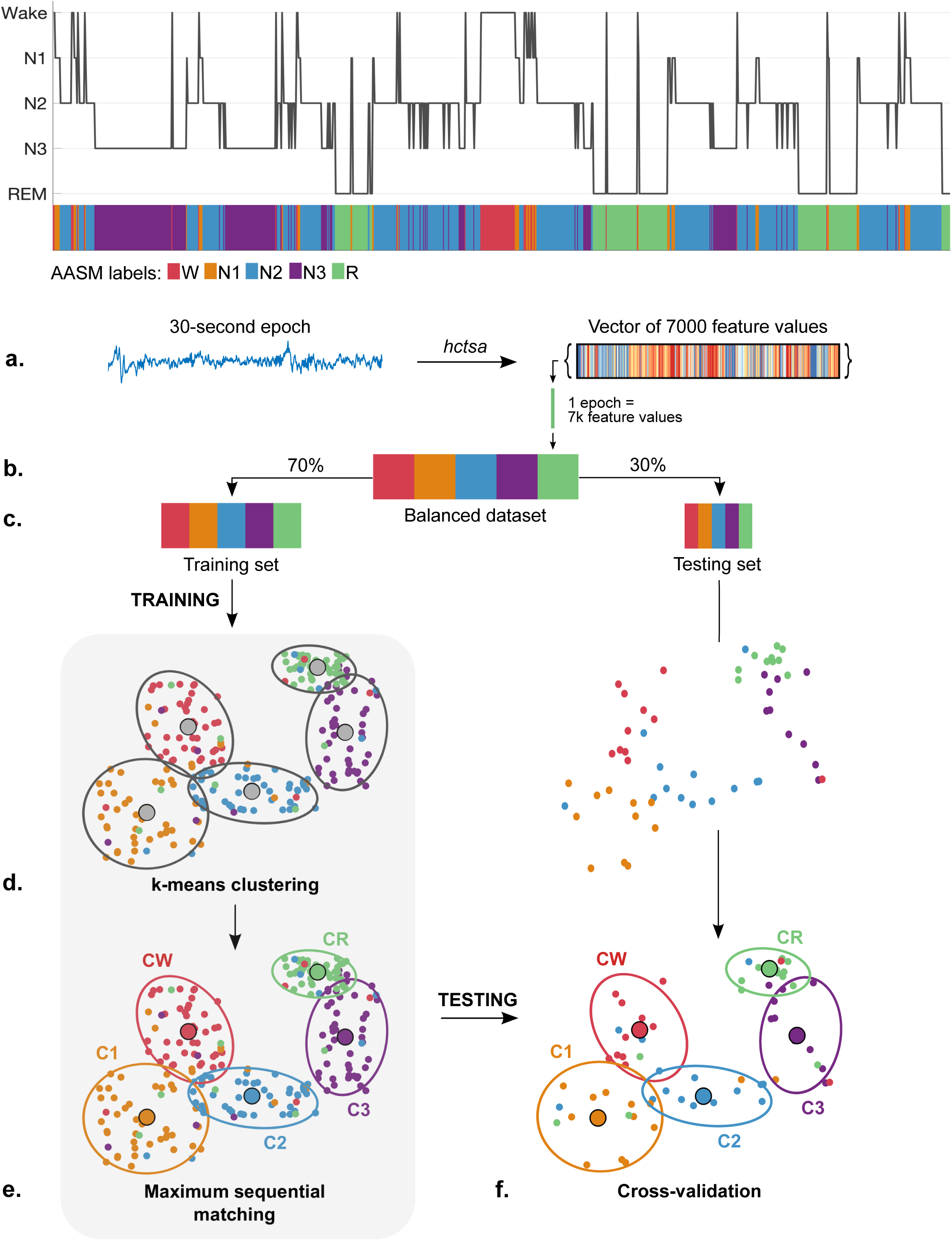
Computational steps in our unsupervised feature-based clustering approach. a) We applied *hctsa* on PSG data, which converted each 30-second epoch into a single vector containing the output values produced by all *hctsa* features for this epoch. b) To equally represent each AASM stage, we formed a balanced dataset by random subsampling. c) We randomly assigned 70% and 30% of the balanced dataset into a training and testing set, respectively. d) We used the feature values of the training set for *k*-means clustering (k=5). Black ellipses depict clusters. Their centroids are depicted as large grey circles. Each small colored dot indicates the given AASM label, representing the vector of feature values from one training epoch. e) Next, we mapped each unlabeled cluster onto one of the five AASM sleep stages based on the most frequently matched AASM label in each cluster using our sequential maximum matching algorithm. We named the 5 clusters as CW, C1, C2, C3 and CR, which were matched with W, N1, N2, N3 and REM respectively. f) After completion of training, each test epoch was assigned to the nearest cluster (i.e., cross-validation). To assess the overlap between cluster decisions and the AASM labels, we computed the percentage of test epochs whose AASM label matches the cluster it was assigned to (e.g., N2 epoch assigned to C2). The entire procedure was iterated 100 times per dataset. See Methods (*Feature-based clustering*) for details on each step of the procedure.

To visualize how the large number of *hctsa* time-series features vary across a night of sleep, we show a color-coded feature matrix representing the feature values for each 30-second EEG, EOG and EMG time series from one subject over one night (Figure 2). The matrix displays the normalized feature values for 5946 valid features on 1014 epochs in chronological order. Below the matrix, we show the hypnogram generated from the AASM labels supplied by consensus between three trained scorers. We observe that the hypnogram generally corresponds with patterns of feature values, notably for the EEG and EOG. More specifically, notice that the short periods of wake (e.g., during the 3rd hour; green arrows at the top) correspond with narrow bands of feature values that contrast with neighboring epochs of N2 sleep. This broad correspondence can be explained by the apparition of high-amplitude hyper-synchronized slow waves at the transition from wakefulness to deep sleep (9, 10), resulting in a sharp increase in the autocorrelation of the EEG signal. The fact that many *hctsa* features reflect these wake-NREM transitions could thus stem from their sensitivity to the signal’s autocorrelation. As a result, the *hctsa* feature matrix broadly tracks the sleep architecture described by visual sleep scoring. However, we also note instances of potential discordances between the two methods. For example, during the continuous REM period (marked by a red bar at the top), multiple patterns of feature values are identifiable. This hints at heterogeneity in time-series properties within conventional sleep stages, which we examine in more detail later.

**Figure 2.**
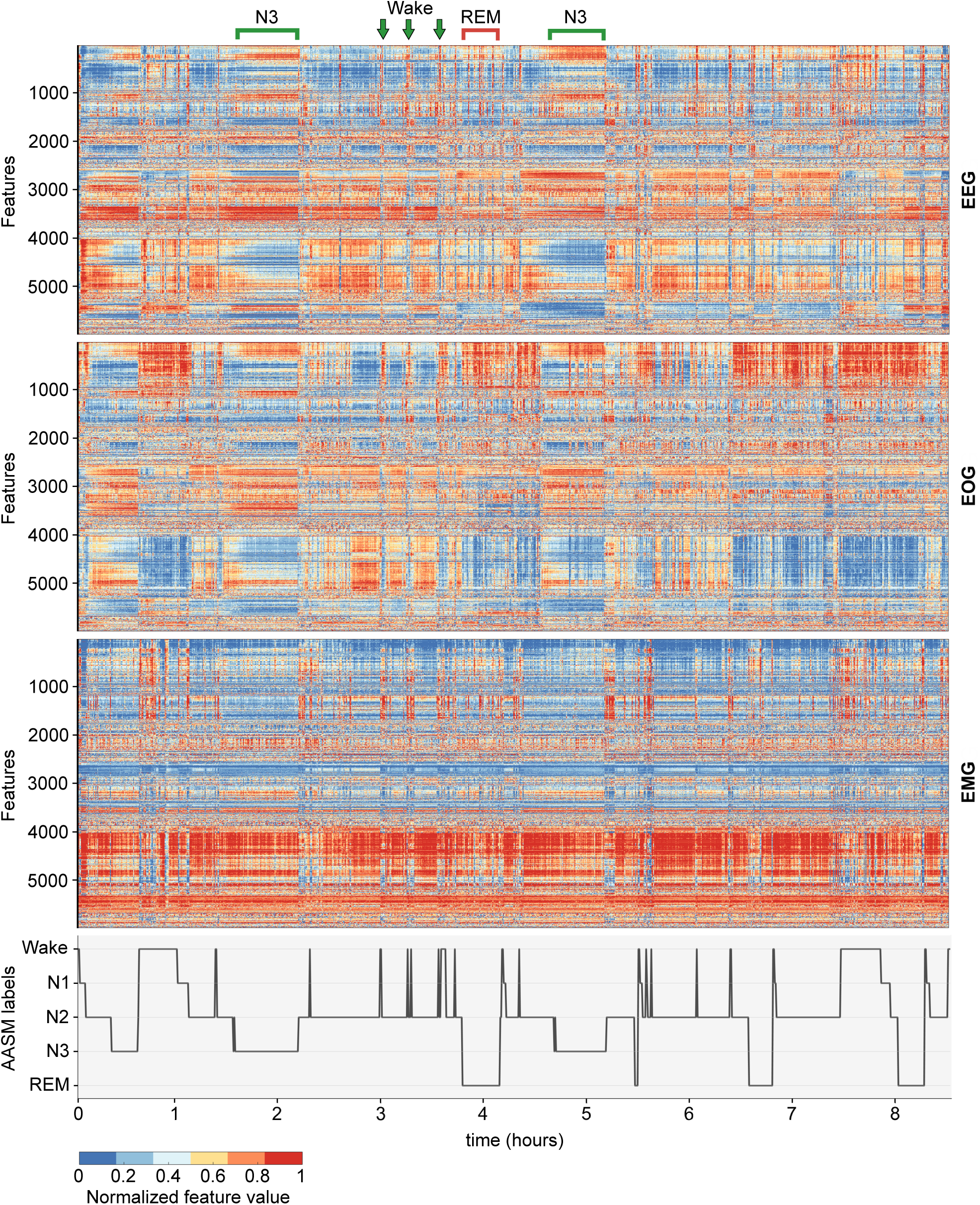
Time-series properties broadly track visual sleep scoring. The matrix is derived from applying 5946 *hctsa* features to one full-night dataset (ID: 1800439). Each column corresponds to one of the 1014 30-second epochs recorded by an EEG, EOG, and EMG (top, middle, and bottom, respectively). The x-axis shows time (in hours) from the first non-wake epoch. Each row corresponds to one *hctsa* feature. To help interpretation, we order features in EEG by their correlation-based similarity using average linkage clustering (18, 19). We order features in EOG and EMG using the same ordering as in EEG. Each pixel encodes a normalized feature value from 0-1 (see colorbar below the matrix). Below the feature matrix, we show the hypnogram (the labels derived by consensus between three trained scorers in accordance with the AASM visual scoring guidelines). The hypnogram is broadly consistent with different patterns in the feature matrix (notably for the EEG and EOG matrices).

### Feature-based clusters substantially overlap with visual sleep scoring

To quantify the degree of correspondence between the two approaches, we computed the mean overlap between the two classifications across datasets (Figure 3a). While the highest overlap was observed for N3 (77.4% of the N3 epochs were assigned to C3), the lowest degree of agreement was obtained for N2 (only 43.1% of the N2 epochs were attributed to C2). The majority of N2 mismatch was due to the assignment into C3 (20.9%). We also note high off-diagonal overlap between Wake, N1, and REM epochs (e.g., 27.6% of N1 assigned to CR, 26.6% of Wake to C1, 21.3% of REM to C1). Overall, this level of overlap demonstrates the ability of our stage mapping algorithm to attribute each cluster to the corresponding AASM stage without *a priori* information on how sleep stages are defined.

**Figure 3.**
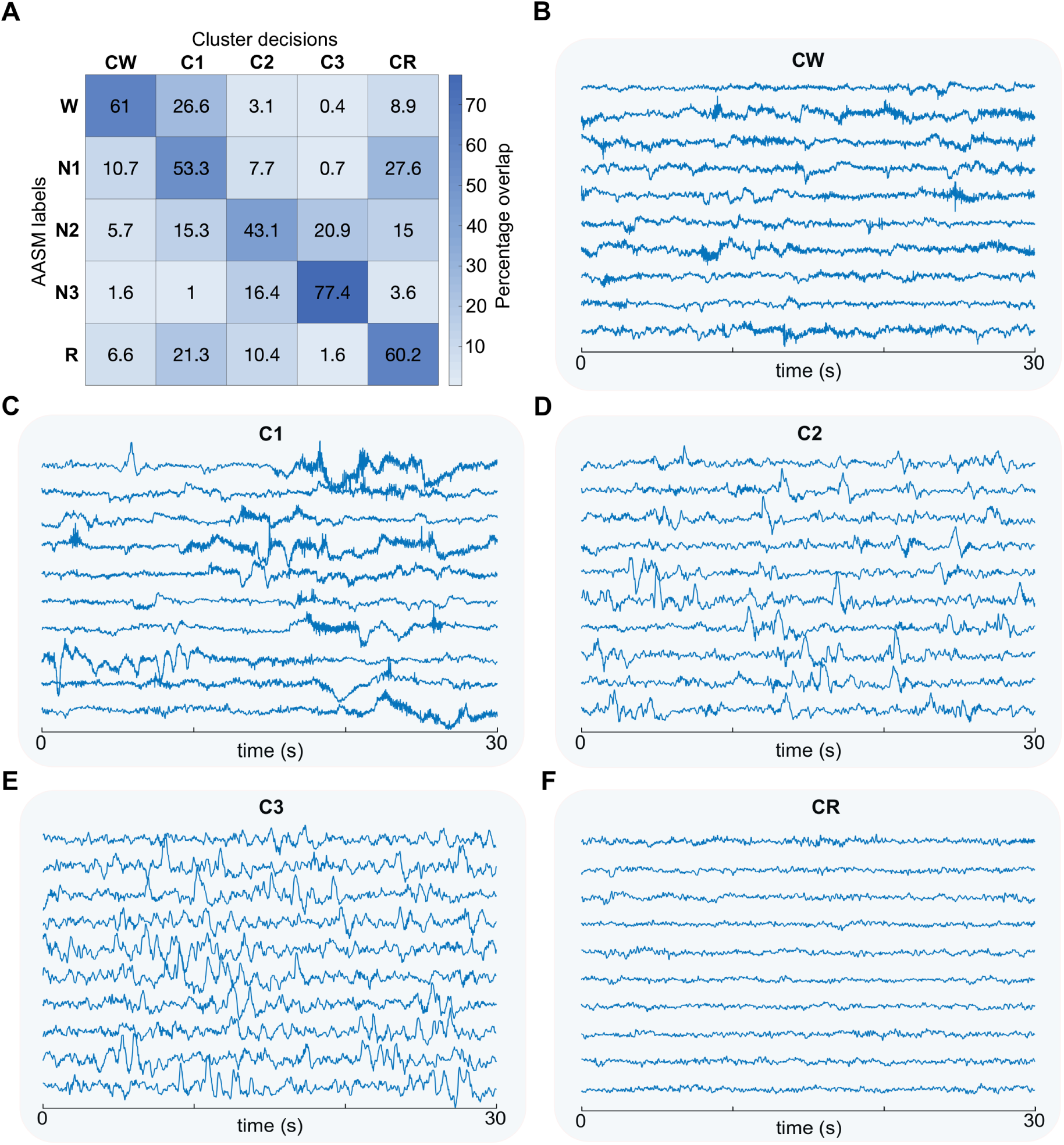
Overlap between the AASM visual scoring and our feature-based clustering across stages. **A**) Overlap matrix between our feature-based clustering results on the test data (columns: CW, C1, C2, C3, and CR) and the AASM labels (rows: Wake, N1, N2, N3, and REM). The number in each cell represents the mean overlap (in %) across the 12 datasets. **B**) CW. **C**) C1. **D**) C2. **E**) C3. **F**) CR. For each cluster, we selected 10 EEG time series (from one dataset; ID: 1800005) that were closest to the cluster centroids. Each line corresponds to the EEG time-series at electrode C3 for one of these 30s-long epochs.

To illustrate what each data-driven cluster looks like, we show exemplar EEG time series located near the cluster centers from one dataset (Figure 3b-f). Feature-based clusters exhibit visual features that are descriptive of AASM sleep stages (e.g., CW and C1 are characterized by high alpha activity, C2 by the presence of K-complexes, C3 by slow-wave activity). This analysis demonstrates that our data-driven approach reveals—without any use of AASM labels—a cluster structure in the data, and this structure reflects the key visual aspects that characterizes traditional scoring.

### Part 2. Revealing the source of divergence between feature-based clustering and traditional scoring

#### Contextual information does not fully explain the observed discrepancies

While the feature-based clusters and traditional sleep-stage labels largely overlap, our novel framework also revealed interesting divergences. Our visual inspection of the epochs with mismatched labels suggested that instances of disagreement may stem from the use of contextual information in AASM scoring. While human scorers are encouraged to use neighboring epochs to assist in labeling ambiguous epochs, our algorithm analyzes each 30-second epoch in isolation.

To test the possibility that the observed divergences are due to the use of contextual information in human scoring, we recruited 4 expert sleep scorers (‘No-context scorers’) and asked them to score a subset of the epochs from the same data and under the same conditions as our feature-based clustering approach. Specifically, we first equated the number of epochs across sleep stages, then presented each epoch in isolation and in random order (see Methods; *Visual scoring task*). Under these settings, no-context scorers agreed with the trained scorers for 67.5% of the epochs. This is lower than the ∼80% interscorer agreement obtained in sleep studies (8), implying that contextual information may explain ∼12.5% of the disagreement. Importantly, no-context scorers agreed with feature-based clustering in 41.1% of cases, similar to the level of agreement with trained scorers (45.2%), suggesting that the use of contextual information does not fully explain the disagreement between feature-based clusters and the implementation of AASM scoring rules.

### Feature-based clustering points to potential sub-stages within AASM sleep stages

To explore the source of discrepancy other than contextual information, in Figure 4 we show the same feature matrix plotted in Figure 2 but with epochs grouped according to the AASM labels (see bottom annotations). Additionally, within each AASM label, we reordered epochs according to the cluster assigned by the algorithm (CW, C1, C2, C3, and CR; see colorbar). Figure 4 shows that, within each AASM stage, epochs assigned to different clusters coincide with visually distinct patterns of EEG and EOG feature values (less so for EMG). This observation is most striking within N2. Feature-based clustering assigned N2 epochs to five different clusters, each displaying distinct patterns in feature values. Figure 5 shows a visualization of the N2 epochs (same data as in Figure 2 and 4) across subjects using *t*-distributed stochastic neighbor embedding (*t*-SNE; see Methods; *t-SNE*). In 10 out of 12 subjects, we observe a heterogeneous structure (marked by dotted lines) within N2. In other words, according to our feature-based clustering approach, N2 is far from being a uniform state.

**Figure 4.**
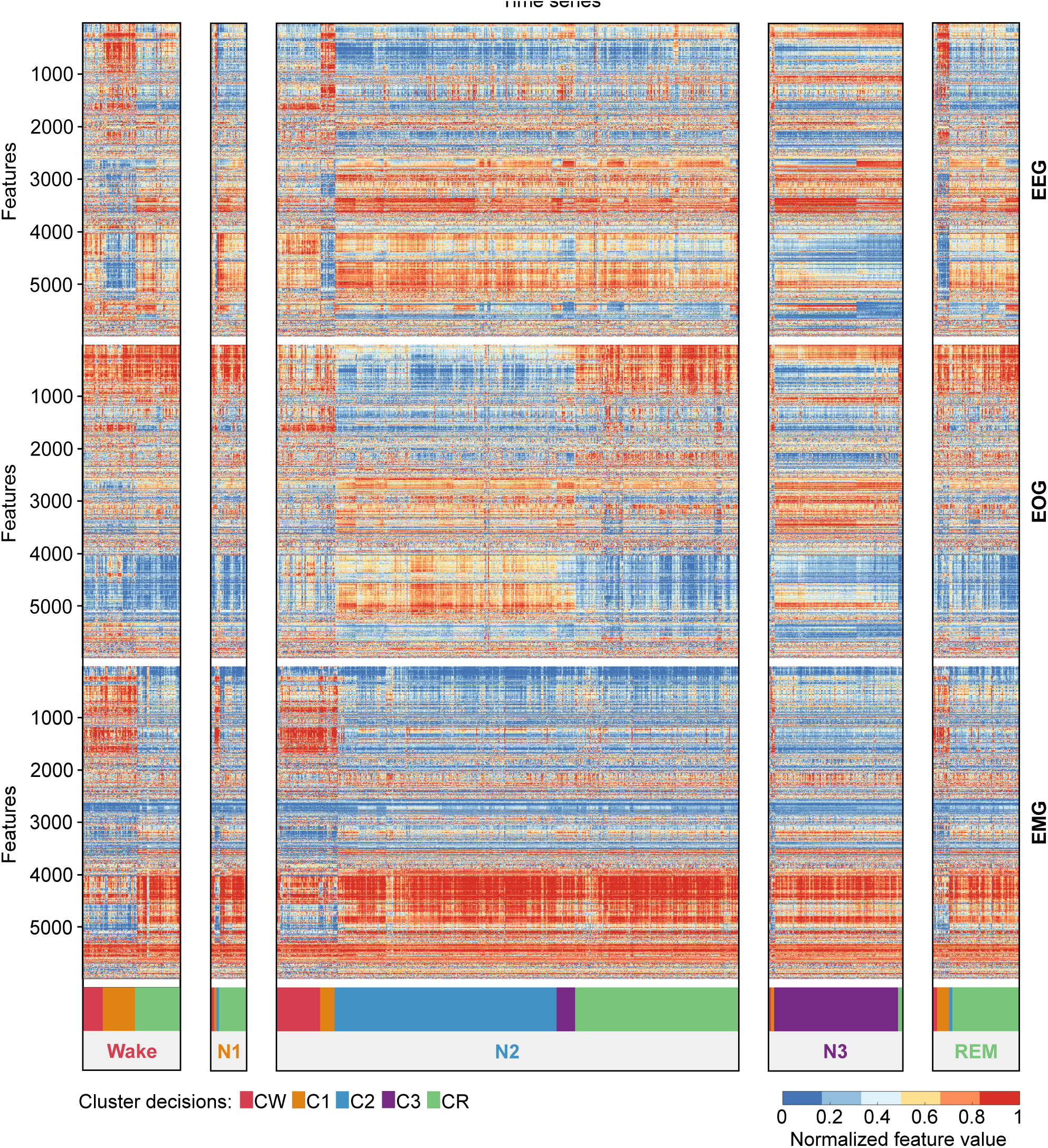
Feature matrix points to heterogeneity within each AASM sleep stage. The same N=1014 epochs in Figure 2 are plotted (see legend of Figure 2), except for a different ordering across the x-axis. From the left to right, we organized columns in five matrices according to the original AASM labels (Wake, N1, N2, N3, REM). Within each AASM label, we further ordered columns according to feature-based cluster assignment (CW, C1, C2, C3, and CR; see colorbar and legend). Disagreements between the AASM labels and feature-based clustering clearly correspond to distinct patterns in the feature matrix.

**Figure 5.**
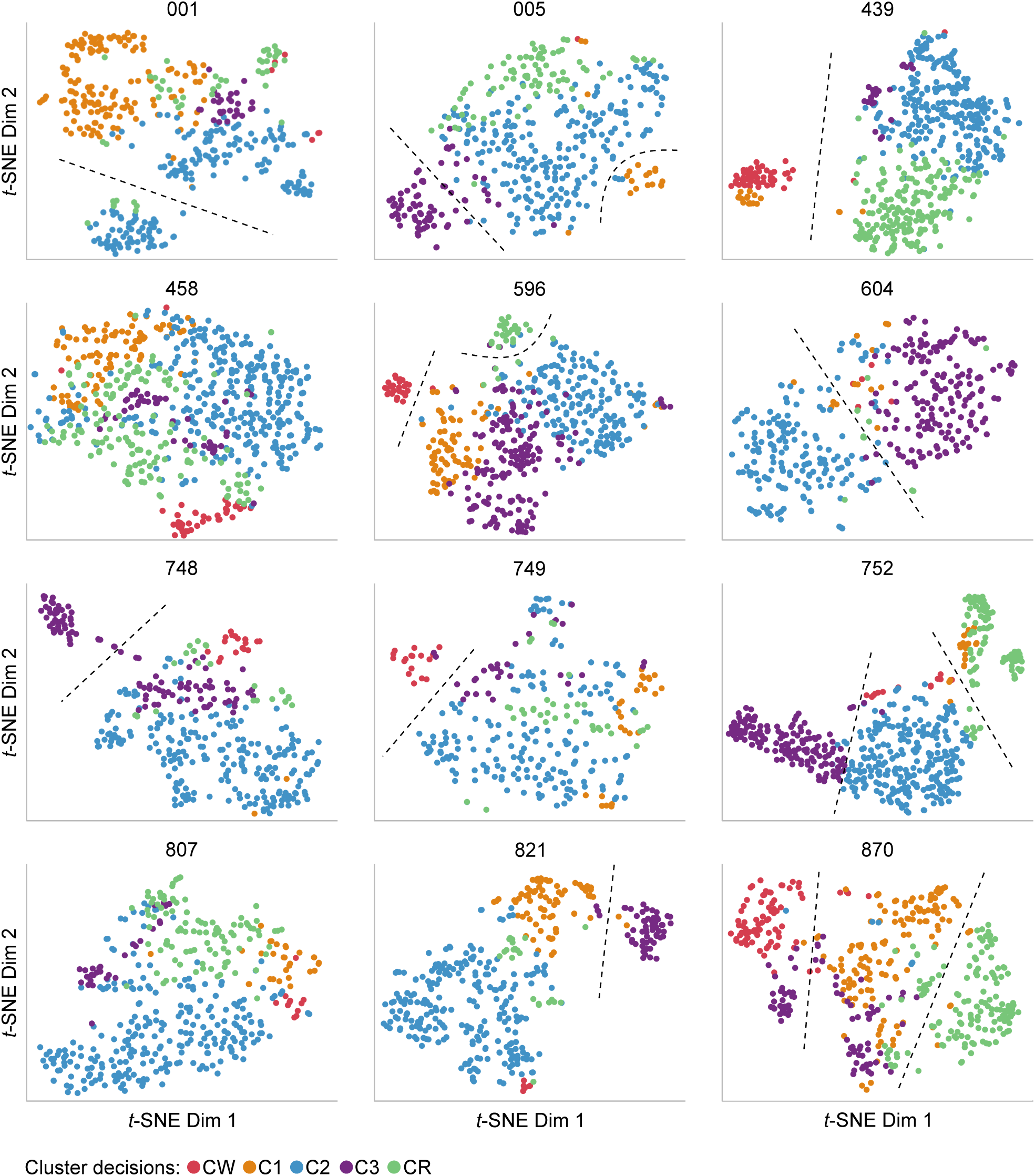
Heterogeneity in time-series properties within N2. *t*-SNE two-dimensional projection of N2 time series (EEG, EOG, and EMG) for each of 12 datasets. Units are arbitrary. The title refers to the dataset index. Cluster assignments (indicated by the color of each dot) were obtained separately for each dataset as in Figure 4 (CW, C1, C2, C3, CR; see legend). For 10 of the 12 datasets, we observe a heterogeneous structure within the N2 stage, as illustrated by black dotted lines.

We conclude that the divergence between feature-based clustering and AASM scoring is driven not just by the lack of contextual information in our approach compared to traditional scoring, but also by the detection of neurophysiological heterogeneity within each AASM stage. The heterogeneity of N2 and other AASM stages has been reported in previous studies and precise characterizations in sub-stages have been proposed (see Discussion). This heterogeneous structure might reflect physiologically distinct periods of sleep that are overlooked with traditional scoring.

### Part 3. Highly discriminative features of AASM sleep stages

In the final part of our paper, we aim to investigate how successfully individual *hctsa* features can discriminate AASM sleep stages. We focus on classifying the four pairs of stages that had the greatest divergence between feature-based clustering and AASM scoring: N1/REM, Wake/N1, N2/N3, and N1/N2 (Figure 3a). This analysis is based on the feature values of EEG time series to identify the most relevant features to the distinction of sleep stages based on minimal PSG information. For this purpose, we now regard the AASM labels as tentative standards against which we quantify the discriminative properties of each single feature. While our ultimate aim is not to replicate traditional scoring, we wanted to investigate whether new types of time-series features, beyond traditional methods for sleep scoring, could be identified using a data-driven analysis with *hctsa*.

### Identification of the most discriminative features

To assess the ability of single time-series features to individually discriminate between a given pair of AASM sleep stages, we performed classification using each *hctsa* feature in isolation (see Methods; *Single-feature analysis*). We computed balanced classification accuracy based on an equal number of epochs in each stage. For each of the four pairs of sleep stages, a large number of *hctsa* features (over 800) were significantly discriminative (*p* < 0.05, FDR-corrected). The maximum classification accuracy across features between N1/REM, Wake/N1, N2/N3 and N1/N2 were 70%, 78%, 91% and 90%, respectively. Given the high performance of individual time-series features, we next aimed to understand the types of time-series properties that these most successful features were measuring in successfully classifying each pair of sleep stages. To achieve this, we inspected the most discriminative features of each sleep-stage pair (including their pairwise dependencies) to infer groups of features that capture similar types of informative differences in sleep-stage dynamics (see Methods; *Single-feature analysis* for details).

For classifying N1 versus REM epochs, the most discriminative time-series features (individual accuracies >65%) measured properties of time-series predictability. A range of features captured these properties, including permutation entropy (20) (up to 69%) automutual information (up to 69%), symbolic motif sequences (up to 68%), and power spectral properties (including the scaling exponent estimated from the power spectrum, 68%). These features indicate that REM time series are more predictable than N1, consistent with the definition of N1 as a variable, transitional state between wake and sleep. We show in Figure 6 the value of the scaling exponent of the power spectrum (a linear fit on logarithmic axes) across EEG time series: N1 has a greater scaling exponent than REM (i.e., a steeper slope of the power spectrum) which suggests a higher level of consciousness in N1, according to recent research in anesthesia, wake and sleep (21, 22).

**Figure 6.**
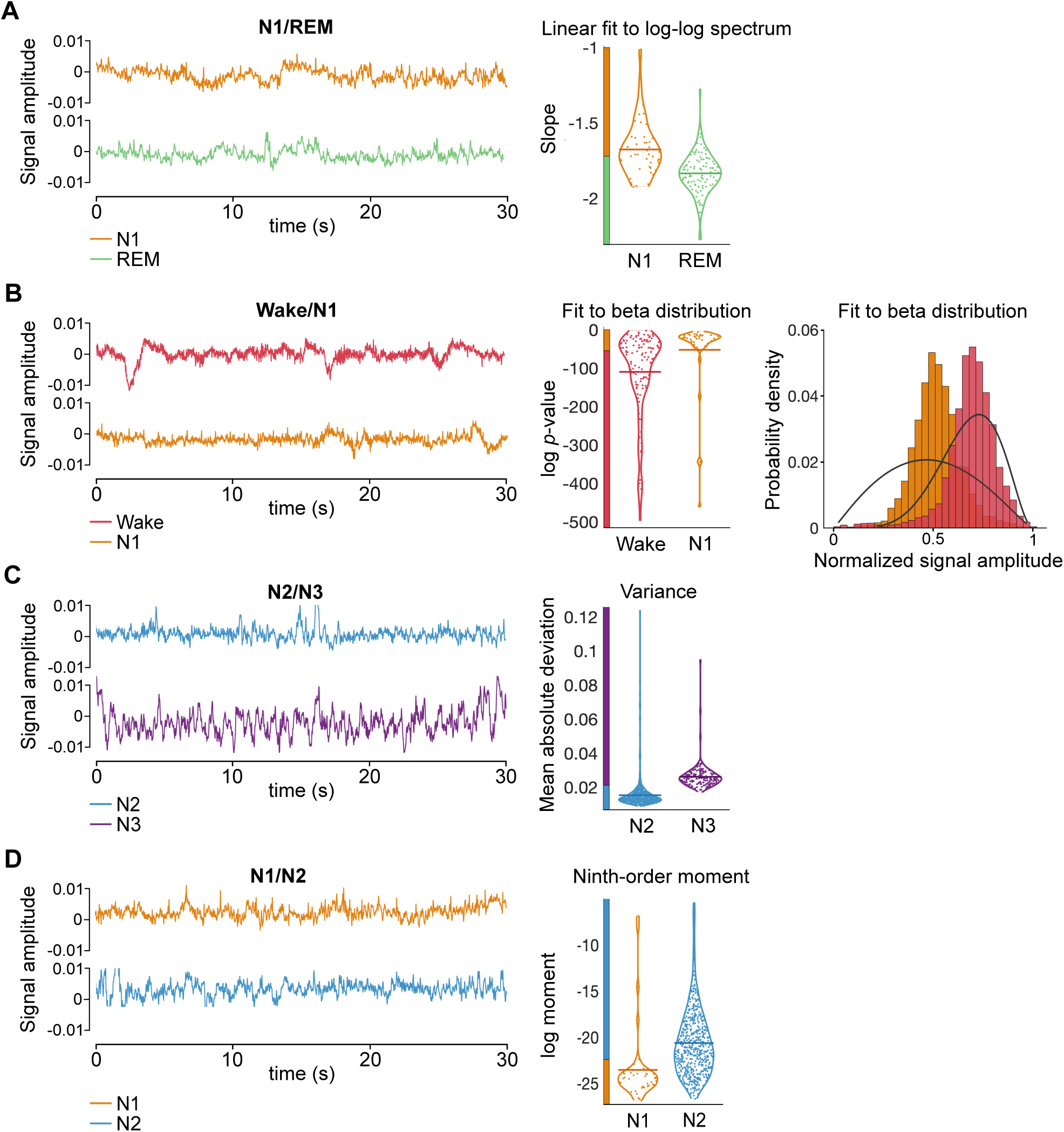
Discriminative features. Discrimination between N1/REM (A), Wake/N1 (B), N2/N3(C) and N1/N2 (D) reached the worst performance (Figure 3a). They were, however, classified rather accurately by the individual features presented here. To understand the discriminative properties of these features, which are rarely used in sleep research, we show the output generated from their application on representative EEG time series from one dataset (ID: 1800001). In the left part, we show the representative EEG time series. In the middle, we show violin plots representing the distribution of feature values generated across EEG time series (represented as dots) in the two stages. The horizontal line corresponds to the mean feature value across time series. For Figure 6B, we provide a visual depiction of the feature output to illustrate how this feature discriminated against Wake and N1. **A**) Slope of the linear fit to the log-log power spectrum (N1/REM). The y-axis of the violin plot shows the slope of the linear fit. **B**) Fit to beta distribution (Wake/N1). The y-axis of the violin plot shows the logarithmic *p*-value generated from the Kolmogorov-Smirnov test—a hypothesis test used to quantify the difference between the beta distribution and the time-series data. On the right, the histograms show the probability density (y-axis) as a function of the normalized EEG signal amplitude (x-axis) from the beta distribution. Black curves depict the fit to the beta distribution. **C**) Variance (N2/N3). The y-axis of the violin plot shows the mean absolute deviation. **D**) Ninth-order distributional moment (N1/N2). The y-axis of the violin plot shows the logarithmic 9th moment.

For Wake and N1, two major groups of features had the highest classification accuracies (>71%): (i) measures of self-correlation and predictability, which indicate that N1 time series are more predictable than Wake; and (ii) measures of distributional outliers, which indicate that Wake dynamics have more outlying data points than N1. The self-correlation features used methods like autoregressive model fits (up to 78%), wavelet decompositions (up to 75%), and Fourier power spectrum properties (e.g., a scaling exponent estimate, 74%). The distributional differences between Wake and N1 could be measured through fits to Beta and Normal distributions, which can capture transient events during Wake such as those depicted in Figure 6B (see left tail of the Wake distribution).

The best performing features for distinguishing N2 and N3 (individual accuracies >85%) measured a range of time-series properties that indicated higher self-correlation (increased predictability) in N3 time series than in N2, and another group of features measuring the increased signal variance in N3 over N2. Highly discriminative self-correlation features include measures of automutual information (up to 90%), local forecasting errors (up to 89%), entropies (up to 87%), and properties of the power spectrum (e.g., scaling exponent estimate, 87%). As for variance, robust increases in EEG amplitudes in N3 relative to N2 were detected by simple features like mean absolute deviation (87%), as shown in Figure 6C. These features characterize N3 time series as more self-correlated than N2, which aligns with the definition of N3 as a state of synchronous activity characterized by high-amplitude slow waves and low signal complexity (23, 24).

Finally, we report a range of time-series features with high individual discrimination of N1 and N2 (accuracies >82%). The first group of features captures the greater number of outliers in N2 epochs, measured using odd distributional moments (e.g., 9th-order distributional moment, 90%) and even distributional moments (e.g., 8th-order moment, 84%). These features could capture the effect of transient physiological events (e.g., K-complexes) that occur during N2, as shown in Figure 6D. A second group of features capture the increased predictability/self-correlation of N2 time series relative to N1, which could be measured by scaling exponent estimates (e.g., a linear fit to the logarithmic power spectrum, 89%, or detrended fluctuation analysis (25) (84%)), or the mean degree of a visibility graph (26) (84%).

Our data-driven approach thus provided interpretable insights into a wide range of time-series properties that differ between sleep stages, and flagged a range of time-series analysis methods that could accurately measure them. The features recapitulated some of the existing sleep-scoring literature (e.g., properties of the Fourier power spectrum, power in some frequency bands, and scaling exponent estimates). But, more importantly for our purposes, they also included a range of novel features that are not widely used to discriminate AASM sleep stages (such as visibility graphs, distributional moments, symbolic motifs and permutation entropies). Incorporating these novel features into automated sleep scoring could provide additional insights into sleep physiology (see Discussion).

## Discussion

We present a novel sleep time-series analysis framework, which aims to go beyond traditional visual sleep scoring. To do so, we 1) leverage over 7700 features to comprehensively characterize sleep time series, 2) cluster time series in an unsupervised way, rather than classifying them according to AASM guidelines, and 3) analyze the data without assuming the temporal context to resolve ambiguities arising in the scoring of sleep-stage transitions. These three steps represent our first attempt to go beyond traditional visual sleep scoring, which leads to a more complete understanding of the underlying neuronal mechanisms of sleep.

In Part 1, we showed that the sleep structure revealed by our approach substantially overlapped with the traditional AASM sleep stages defined by visual scoring (Figure 2), suggesting that both approaches capture a genuine physiological structure of sleep. However, we also observed important points of divergence between the two methods (Figure 3a). In Part 2, we examined potential sources of the divergence. While it is partially explained by contextual information, it is also reflected by the fact that *hctsa* features depicted heterogeneity in time-series properties within each of the AASM sleep stages (Figure 4 & 5). This promising result is further followed up in Part 3, where we examined the discriminative properties of high-performing features (Figure 6). These features promise novel discoveries of the structure of sleep in future studies adopting our framework. In the rest of Discussion, we identify various directions to which our framework can readily extend to pick up low-hanging fruits in its applications.

### Towards establishing a general and objective ground truth of sleep stages

We used AASM standards as a benchmark to examine how our framework characterized and organized sleep. Although AASM visual scoring is the gold standard for sleep staging, we do not consider this to be the ground truth of sleep physiology. As emphasized in the Introduction, we argue that conventional visual scoring provides a rather inadequate description of the physiological changes that occur during sleep. Given the lack of ground truth, our study raises the question of what a better sleep classification approach would involve and, more specifically, what “better” means in this context. Our data-driven approach represents a highly flexible framework which can adapt to different definitions of sleep and sleep stages. The high number of features offers the possibility to fine-tune a wide range of parameters and therefore to adjust to various cases in which sleep needs to be assessed. This includes, for example, selecting features that are the most sensitive to a specific sleep stage, sleep function or sleep pathology. In this regard, our approach has significant potential to contribute to a more complete understanding of the nature of sleep.

One possible future research direction is to apply our framework to data from the same subjects across multiple nights. By dividing the data into training and test nights, we can apply similar cross-validations as we established in our framework (Figure 1). Such studies would be an ideal way to estimate the sensitivity and specificity of our framework, providing a benchmark for further studies.

The confirmation of the stability and precision of our framework, in comparison with the traditional AASM scoring approach, can also be tested across subjects. How much individual variability will our framework reveal on its application to massive healthy population data? Can it explain variance in mental and physiological differences in a more sensitive and specific way than traditional approaches? While the AASM guidelines are explicit in not applying the same classification rules to non-normative data (e.g., patients with sleep disorders, infants or seniors), our framework has no such inherent limitations. Broader applications of our framework can eventually establish a new norm and potentially replace the current gold standard of sleep scoring.

### Refining traditional sleep stages

In Part 2, we took an initial step towards the discovery of sub-stages that are overlooked with the traditional scoring. We focused on N2 because this stage had the lowest level of agreement between our approach and AASM scoring (43.1%; Figure 3a) and represented the largest amount of the data available in each dataset (Figure 4). t-SNE projection exhibited heterogeneity within N2 in a visually striking way and in the majority of datasets (Figure 5).

Variability within N2 was previously reported. Sleep spindle activity does not occur uniformly within N2, but their occurrence is modulated across and within sleep cycles (27) and is dependent on local temporal context (28), which may reflect fluctuations in sleep depth within N2 (29). Along this line, based on autonomic and hormonal activity, the division of N2 has been proposed as a quiet type (before the transition into N3) and an active type (preceding REM sleep) (30). From a physiological perspective, it is not surprising if N2, which occupies as much as 50% of total sleep (31), is not physiologically homogeneous.

While we focused on N2 in Figure 5, we believe that this observation is not restricted to this stage. The feature matrix presented in Figure 4 suggests variability in time-series properties within every AASM sleep stage. To the extent that these variations reflect underlying changes of sleep, our findings indicate that sleep staging can be refined to unveil more subtle sleep stage distinctions. These distinctions may be defined within AASM sleep stages, similar to how REM sleep can be decomposed into a phasic and tonic period (32), each of which is associated with different physiological and cognitive processes (33). Sub-stages may also not necessarily align with coarse sleep stages but instead emerge as transitional stages, similar to how N1 briefly appears between the wake state and N2.

The delineation of sub-stages can also be achieved using much shorter time scales than the traditional 30-second time window. As the present study sought to compare the structure emerging from feature-based clustering with that of traditional scoring, we adopted the 30-second window as the unit of our analysis. A previous study showed that the AASM scoring can be successfully replicated using time windows down to 5s (34). Our framework is highly flexible too and can be applied at shorter time scales, which is likely to be optimal for the reliable identification of sub-stages, as demonstrated for example within N1 (35). The temporal refinement is also most suited to the incorporation of short-lived events, such as sleep spindles or K-complexes, which may be associated with the ability to report dream content (36, 37). Application of our automated framework therefore promises highly productive areas of future research by refining the temporal scales of the analysis.

Our approach is also scalable in space. In fact, the EEG technology (as well as other physiological variables, other than EOG and EMG) has progressed tremendously in quality and quantity since the time of the original development of the AASM guidelines. Especially, the high quality and high-density EEG (60-256 channels) are more and more adopted in sleep research in both healthy and clinical populations. Higher spatial resolution in EEG has led to discoveries that have reshaped our conceptualization of sleep physiology, with the characterization of NREM slow waves as travelling waves (38), the identification of two types of NREM slow waves (39), the presence of slow waves in REM sleep (40) and even in wakefulness (41), or the identification of inter-hemispheric differences in slow wave activity (42). This recent research has revealed the importance of the local aspects of sleep (43). This notion of “local sleep” challenges the view that sleep is a global phenomenon and raises the question of locally defined sub-stages (44). While these studies point to the importance of analyzing all available channels of EEG, the AASM guidelines ignore much of the spatial information. The incorporation of spatial information in our framework is highly feasible and represents another low-hanging fruit for its application.

### Broadening the characterization of sleep time series

To our knowledge, a comprehensive and interdisciplinary library of time-series analysis, such as *highly comparative time-series analysis* (*hctsa* (18, 19)), has never been systematically applied to sleep EEG data. While sleep scoring studies typically investigate the significance of single measures, *hctsa* offers the opportunity to assess the discriminative power of over 7700 features at once. In the last part of the paper, we presented highly discriminative features for the distinction of four pairs of stages (N1/REM, Wake/N1, N2/N3 and N1/N2). These features could incorporate additional meaningful information in sleep scoring and refine the current description of sleep architecture.

While some may be concerned that our approach will drown the researchers in the massive number of features in space and time, this problem can now be overcome by leveraging machine learning techniques (34), a highly promising interdisciplinary research agenda. Employing automated machine learning, recent studies aimed to expedite and replicate traditional sleep scoring by humans (see 45 for a review). While this is not what we aimed for in the present work, we see a similar approach can be usefully incorporated within our framework.

Our data-driven approach recapitulated some of the existing sleep scoring literature. Among the most discriminative features, we found measures that are traditionally used (i.e., power in specific frequency bands). We also identified less frequently used features that have also been described as discriminative in past automatic scoring studies, such as measures of wavelet transform (46) and permutation entropy (47). Another interesting feature highlighted by our analyses is the slope of the linear fit to the power spectrum—the most discriminative feature across all 10 pairs of stages (not reported in Results)—which has been recently used to track changes in vigilance state (22) or consciousness (21).

Importantly, we also found highly performing features that have rarely received attention, including visibility graphs, symbolic motifs, automutual information, distributional shape and multiscale entropy. Interestingly, most of these features probe nonlinear properties of the signal. Given the nonlinear complexity of biological signals, these features are more suited to assessing the intrinsic dynamics of EEG during sleep than Fourier-based spectral analysis, which assumes linearity and stationarity of the signal (48). Furthermore, nonlinear features are of high clinical value for characterizing sleep physiology in ageing and sleep disorders, where brain dynamics are altered (49). Our data-driven analysis points to the importance of using such features to capture the complex dynamics of sleep.

Our approach examined all of these features at once without cherry-picking which features to test and report. Selective testing and reporting of a single feature in an individual study runs a significant risk in the replication crisis. This is a pervasive problem across the scientific fields, but in particular, it is a significant problem in the young field of data-analysis intense neuroscience (e.g., 50). Such risk can be minimized by our approach using *hctsa*.

### Future implications

As we have already mentioned, there are already various feasible and impactful research projects based on our framework. As such, this paper is a first step towards the establishment of a purely data-driven sleep classification method to discover sleep sub-stages that are not visible to the human eye. We also note that further qualitatively distinct development can be achieved by extending our framework here with multivariate analysis (under preparation, Cliff, Tsuchiya, Lizier, Fulcher). Multivariate analysis, such as correlation and coherence, has been attracting attention in signal processing of the task-related EEG analysis, but rarely utilized in sleep research. Systematic investigations of temporally lagged correlations between electrodes are likely to characterize functional and effective connectivity among brain areas, which would arrive at a more reliable and fine-grained definition of sleep stages based on brain mechanisms.

In terms of application of our framework, we believe that clinical applications are highly promising. The AASM visual scoring rules were originally developed for healthy subjects. Conventional scoring may be too coarse to capture relevant structural patterns in patients with sleep disorders, whose sleep is often fragmented (51). Moreover, sleep architecture is generally characterized by an abnormal time course of sleep stages in sleep disorders (52), therefore the use of contextual information is less adequate for the classification of transitional epochs in patients. Our framework would form the basis of an objective method to better classify sleep states of these patients that have been outside of the purview of the traditional sleep scoring.

Furthermore, our framework could be used as a complement to existing scoring conventions in order to reach a more precise understanding of the PSG correlates of a certain sleep state or phenomenon. In sleep state misperception or paradoxical insomnia, someone’s conventionally scored PSG recording indicates that they are asleep or display normal sleep patterns, while they report being awake or having barely slept (53, 54). Not only does current PSG analysis appear not to reflect substantial aspects of the biological process of sleep, but it also correlates poorly with subjective ratings of sleep quality (55). However, more in-depth analysis of PSG data may allow us to reconcile the subjective reports with PSG features (56). *hctsa* could therefore be a useful tool for determining how PSG data during sleep state misperception differs from normally perceived sleep (57).

Another large area of application is to find neural signatures of various conscious states. Dream research, for example, can contribute to the theoretical understanding of consciousness (58). However, dreaming does not map onto one specific AASM stage (59-61). As revealed in Part 3, there are features available in *hctsa*, such as the linear fit to the logarithmic power spectrum, that have the potential to act as reliable brain signatures for conscious experience during sleep. It is highly surprising and enigmatic if one of the most vivid and distinct experiences in our life, dreaming, does not have a clear neural basis.

In fact, our framework opens a new direction of consciousness research: the data-driven search for conscious states. For example, by identifying distinct feature patterns in ongoing EEG, sleeping participants can be awakened and asked to report their conscious experience (62, 63). Likewise, wakeful subjects can be asked to report their ongoing thoughts (41, 67). This reversal of analysis from brain to phenomenology is technically more demanding, but it is one of the holy grails of neuroscience. Our framework complements the standard approach that starts from phenomenology or from functional considerations and behavior to categorize conscious states. Ultimately, these different approaches can be unified into a single research program.

Our novel framework, which applied a massive feature extraction tool (*hctsa*) to sleep data, promises a pathway to go beyond traditional sleep staging, which has been outdated since its original proposal. While our framework provided broadly consistent classification results, it revealed finer structures than the traditional approach. Once validated with more extensive within- and across-participants validation projects, our framework can potentially establish a new norm, which generalizes across diverse populations. It can further help understand one of the biggest remaining mysteries in current science: the neural basis of conscious experience.

## Methods

### Feature-based clustering

#### Pre-processing and selection of CCSHS data

We selected full-night polysomnographic recordings from 12 participants (6 female & 6 male) from an open-source dataset from the Cleveland Children’s Sleep and Health Study (CCSHS) (65-69). The dataset contains recordings from adolescents aged 16 to 19 years and is available for download at https://www.sleepdata.org. Each recording contained electroencephalograms (C3/C2 and C4/C1, A1, & A2), bilateral electrooculograms, a bipolar submental electromyogram, sampled at 256Hz for the EMG and 128Hz for the EEG. We selected 1 EOG derivation (LOC-A2: electrode placed next to the left canthus and referenced to the opposite mastoid, A2), 1 EMG derivation (EMG1-EMG2, bipolar derivation over the chin) and 1 left central scalp EEG derivation reference to the opposite (right) mastoid (C3-A2). This central EEG derivation is optimal for the detection of both sleep spindles and slow-waves. To ensure a consistent sampling rate across the three selected channels, we downsampled the EMG channel from 256 Hz to 128 Hz. We used MATLAB (R2015b) to process CCSHS data.

### Feature extraction and selection

We performed feature extraction on MATLAB 2015b using *hctsa* v0.95. This version includes 7749 features that are derived from diverse scientific methodological literatures (such as physics, seismology, economics, neuroscience, and artificially generated simulated data). Among the 7749 *hctsa* features, some returned special values (non-real values or fatal errors). We excluded these features using the *hctsa* function *TS_Normalise* (19). Using this function, we removed the features that generated special values in more than 20% of the time series of each dataset. On average, we removed 1747 features per dataset (min=1174, max=1981).

The range of values generated by the remaining *hctsa* features varied across features. Therefore, we normalized values using the Scaled Robust Sigmoid (SRS) transformation implemented in *TS_Normalise*. SRS is a nonlinear transformation that uses median and interquartile ranges instead of the mean and standard deviation. SRS is robust against the influence of outliers and it scales the transformed data into a range between 0 and 1 (negative infinity maps to 0 and positive infinity maps to 1). The normalization by SRS of each feature value *x* can be defined as:

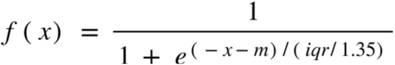

Where *m* is the median value produced by the given feature across time series and *iqr* is the interquartile range across time series.

### Balanced datasets

Based on the AASM labels provided by the trained scorers, we trimmed the 12 selected datasets (Table S1) to exclude any periods of wakefulness that occurred directly before sleep onset (defined as the first non-wake epoch) or directly after the final non-wake epoch. This allowed us to focus on the intra-sleep periods of wakefulness.

The five sleep stages did not occur in equal proportions throughout the night. Table S1 gives the proportion of each AASM stage, which is similar to the known proportion of sleep states (e.g., N1 comprises only 5% of sleep whereas N2 accounts for approximately 50% (31)). To deal with the imbalance, we subsampled the datasets to equalize the number of epochs for each stage as explained in Figure 1b. We took all of the epochs (*N*) of the least frequent stage for each dataset, then randomly selected *N* epochs from each of the other five stages (total 5*N* epochs). We have selected the datasets with *N* ≥ 37 (minimum total epochs = 185). This process underwent 100 iterations, at each of which new epochs were randomly selected from the unbalanced dataset to form the balanced dataset.

### k-means clustering

We used *k*-means clustering (MATLAB function *kmeans* with Replicates = 50, Max Iterations = 500 from Statistics and Machine Learning Toolbox version 11.2). This algorithm groups data based on its Euclidean distance to the center location for each of *k* clusters (70). We used it due to its simplicity, efficiency and wide presence in the literature (71). To enable comparison with the 5-way classification of the AASM guidelines, we set k to be 5. However, by no means does our framework impose this number.

### Sequential maximum matching

When comparing the unsupervised clustering results with the AASM labeling, we assigned each cluster into one of the 5 AASM sleep stages (Figure 1e), according to an algorithm we call “sequential maximum matching”. This matches each cluster to one of the five AASM sleep stages based on the most represented AASM label in the cluster. For details, see Figure S1. When there was a tie, we selected a cluster according to the cluster order randomly generated by the k-means algorithm.

### Visual scoring task

During standard visual scoring, human scorers have access to a full night of temporally sequenced data. This allows them to rely on temporal context to score the epoch, especially when it is ambiguous (e.g., is it preceded/followed by N2, N3 or REM?). To compare the epoch-by-epoch discrimination in our approach with traditional scoring, we created a task where the surrounding context was eliminated for human scorers. We recruited 4 scorers (‘No-context scorers’), who had substantial experience in sleep scoring according to the AASM guidelines (all from the sleep laboratory at Monash University including one of the authors, TA). They scored isolated 30-second epochs one at a time in a 5-way forced choice manner using our custom-made sleep scoring experimental software (MATLAB, Psychtoolbox). In each trial, the software displayed an isolated 30-second signal consisting of 1 EEG (C3-A2), EOG (LOC-A2) and EMG (EMG1-EMG2). Participants were asked to score these epochs in accordance with the AASM sleep scoring guidelines and to the best of their ability. The number of epochs for each sleep stage was equalized. Two participants scored a subset of randomly selected epochs (n=37 trials per stage, in total n=185) from a balanced dataset (ID: 1800001) and the two others scored epochs (n=41 trials per stage, in total n=205) from a different balanced dataset (ID: 1800005).

### t-SNE

To provide a visualization of N2 time series from the *hctsa* feature space, we used *t* -distributed stochastic neighbor embedding (*t*-SNE), which projects high-dimensional data into a low-dimensional space (72). We performed a two-dimensional *t*-SNE projection with MATLAB’s function *tsne* (default parameters) using as input the EEG, EOG and EMG feature values of the N2 time series.

### Single-feature analysis

In Part 3, we assessed the ability of single *hctsa* features to individually discriminate between N1/REM, Wake/N1, N2/N3 and N1/N2. In this analysis, we focused on the feature values generated by the EEG derivation. Given that some *hctsa* features produced special values in some datasets and not others, we filtered out such features in any of the 12 datasets (2146 non-valid features) to enable the combination of results across datasets. A total of 5603 valid features that were common to all datasets were considered for single-feature analysis.

### Computation of classification accuracy

To assess the discriminative power of *hctsa* features, we calculated the cross-validated classification accuracy generated by each feature individually. We selected an equal number N of epochs in each pair (N corresponds to the number of epochs from the least frequent stage) to obtain a balanced accuracy across classes. We performed cross-validation using the leave-one-out strategy: we selected N-1 epochs of each stage as the training set and the left-out epoch as the testing set. Here, we followed a median-split method to assign individual data points (i.e., 1 EEG feature value from a test epoch) to one of the two stages. First, we calculated the median of the feature values of the set of training epochs for each of the two stages. We then calculated the threshold, which is the middle point between the two medians. Each test epoch was then assigned to one of the two stages depending on whether its feature value was higher or lower than the threshold. We repeated leave-one-out cross-validation N times to allow every epoch to be used for testing. We averaged the percentage balanced accuracy generated by each feature across 100 iterations.

### Computation of *p*-values

To determine which features were significantly informative of sleep stages, we performed permutation testing by randomly shuffling the AASM labels of EEG time series. We computed balanced classification accuracy (following the procedure described in *Computation of classification accuracy*) over 100 iterations based on permuted labels, producing for each given feature and dataset a null distribution of 100 samples. Each sample was averaged across datasets. We estimated *p*-values as the proportion of null samples whose mean accuracy was as high or higher than the mean accuracy generated from un-permuted labels. We corrected *p*-values using the False Discovery Rate (FDR) method (73).

### Pairwise dependency

To assess the dependency between the most discriminative features, we computed the pairwise dependency (Spearman’s correlation) between the 40 highest performing features of each pair of stages to determine which measures captured unique concepts of the EEG signal. We considered features as highly intercorrelated if their absolute pairwise correlation coefficient (|R|) was >0.8.

## Acknowledgements

We thank Tom Drummond for co-supervising PS for her thesis, Dror Cohen for his pilot analysis using the *hctsa* method on another set of sleep data, Jeroen van Boxtel for discussion on the visual scoring task, Jay Kim for constructing a preliminary version of the visual scoring task, Romain Holzmann and Ursula Voss for providing the data, whose results will be published elsewhere.

## Funding

National Health and Medical Research Council (APP1183280) (NT, TA, JMW).

National Health and Medical Research Council (GNT1183280) (BDF).

Australian Research Council (DP180104128 and DP180100396) and Grant-in-Aid for Transformative Research Areas (B) (HDM5Y) from Japan Society for the Promotion of Science (NT).

International Brain Research Organization Post-Doc fellowship and Long-Term Fellowship from the Human Frontier Science Program (LT000362/2018-L) (TA).

## Author contribution

Conceptualization: NT, JMW

Methodology: PS, NT, JMW

Investigation: ND, ZHK, JW

Supervision: NT, TA, BDF

Writing: ND, JW, ZH

## Data and materials availability

All data needed to evaluate the conclusions in the paper are present in the paper and/or the Supplementary Materials. Additional data related to this paper may be requested from the authors.

