## Supplementary Table 1, Supplementary Figure 1 for "Beyond traditional visual sleep scoring: massive feature extraction and unsupervised clustering of sleep time series"

### Supplementary Materials

| Dataset number | Dataset ID | Gender | Age | Wake (%) | N1 (%) | N2 (%) | N3 (%) | REM (%) | Duration (min) |
|---|---|---|---|---|---|---|---|---|---|
| 1 | 1800001 | F | 17.2 | 9.02 | 3.44 | 38.91 | 26.18 | 22.45 | 490 |
| 2 | 1800005 | M | 17.3 | 7.24 | 3.79 | 39.34 | 28.48 | 21.15 | 501 |
| 3 | 1800439 | M | 17.7 | 12.04 | 4.34 | 56.66 | 16.39 | 10.56 | 530 |
| 4 | 1800458 | F | 18.9 | 6.02 | 5.29 | 58.85 | 12.96 | 16.88 | 548 |
| 5 | 1800596 | F | 17.8 | 11.96 | 3.99 | 44.91 | 21.70 | 17.45 | 565 |
| 6 | 1800604 | F | 17.7 | 12.26 | 4.61 | 39.10 | 21.38 | 22.64 | 539 |
| 7 | 1800748 | F | 17.7 | 34.33 | 5.21 | 37.34 | 14.71 | 8.41 | 468 |
| 8 | 1800749 | M | 17.4 | 5.22 | 4.47 | 30.56 | 41.32 | 18.42 | 504 |
| 9 | 1800752 | M | 17.7 | 9.39 | 5.49 | 58.97 | 6.86 | 19.30 | 493 |
| 10 | 1800807 | M | 18.2 | 10.86 | 4.66 | 47.67 | 22.06 | 14.75 | 527 |
| 11 | 1800821 | F | 17.7 | 17.68 | 4.40 | 40.56 | 17.98 | 19.38 | 539 |
| 12 | 1800870 | M | 17.2 | 16.15 | 5.28 | 44.07 | 22.23 | 12.26 | 513 |
| | | **Average** | **17.7** | **12.68** | **4.58** | **44.75** | **21.02** | **16.01** | **518** |

**Table S1. Participant sleep metrics.** Participant sleep metrics for the 12 full-night recordings (6 male and 6 female participants selected at random) before trimming and balancing datasets. Columns 5-9 show the percentage of time spent in each of the five AASM sleep stages (Wake, N1, N2, N3, REM) as defined by consensus between the three trained scorers. The bottom row shows the average percentage and duration of the data across the 12 datasets.

AASM labels

| *k*-means clusters | W | N1 | N2 | N3 | REM | Order of selection | Assigned cluster |
|---|---|---|---|---|---|---|---|
| **Cluster 1** | 0 | 0 | 1 | 38 | 0 | 1st | **C3** |
| **Cluster 2** | 22 | 5 | 6 | 3 | 4 | 3rd | **CW** |
| **Cluster 3** | 8 | 17 | 11 | 1 | 27 | 2nd | **CR** |
| **Cluster 4** | 0 | 1 | 22 | 0 | 3 | 4th | **C2** |
| **Cluster 5** | 12 | 19 | 2 | 0 | 8 | 5th | **C1** |

**Figure S1. Sequential maximum mapping**. We illustrate the procedure used to map each *k*-means cluster (rows) to one of the five AASM sleep stages based on the number of AASM labels (columns) in these clusters, using representative fictitious data (i.e., a balanced dataset with 42 epochs per sleep stage). The algorithm begins by selecting the cluster that has the highest number of epochs from one of the 5 AASM stages. Here, Cluster 1 is selected first as it contains the highest number of epochs from N3 (N=38; see column 'Order of selection'). Because this high number of epochs belong to N3, Cluster 1 is assigned to C3. The algorithm then selects the next highest value and maps the cluster to the corresponding stage. In this example, Cluster 3 contains the second highest number of epochs (N=27) labeled as REM, and is therefore assigned to CR. When a tie-break is necessary (where two clusters both have the next highest number of epochs belonging to one stage), the cluster is selected according to cluster order (in this case, Cluster 2 is selected before Cluster 4). The process stops when all 5 *k*-means clusters are mapped onto one of the 5 stages.
